# LRRK2 G2019S promotes the development of colon cancer via modulating intestinal inflammation

**DOI:** 10.1101/2023.06.28.546897

**Authors:** Yuhang Wang, Joyce Z Gao, Taylor Sakaguchi, Thorsten Maretzky, Prajwal Gurung, Sarah Short, Yiqin Xiong, Zizhen Kang

## Abstract

LRRK2 G2019S is the most prevalent variant associated with Parkinson’s disease (PD), found in 1-3% of sporadic and 4-8% of familial PD cases. Intriguingly, emerging clinical studies have suggested that LRRK2 G2019S carriers have an increased risk of cancers including colorectal cancer. However, the underlying mechanisms of the positive correlation between LRRK2-G2019S and colorectal cancer remain unknown. Using a mouse model of colitis-associated cancer (CAC) and LRRK2 G2019S knockin (KI) mice, here we report that LRRK2 G2019S promotes the pathogenesis of colon cancer as evidenced by increased tumor number and tumor size in LRRK2 G2019S KI mice. LRRK2 G2019S promoted intestinal epithelial cell proliferation and inflammation within the tumor microenvironment. Mechanistically, we found that LRRK2 G2019S KI mice are more susceptible to dextran sulfate sodium (DSS)-induced colitis. Suppressing the kinase activity of LRRK2 ameliorated the severity of colitis in both LRRK2 G2019S KI and WT mice. At the molecular level, our investigation unveiled that LRRK2 G2019S promotes the production of reactive oxygen species, triggers inflammasome activation, and induces cell necrosis in the gut epithelium in a mouse model of colitis. Collectively, our data provide direct evidence that gain-of-kinase activity in LRRK2 promotes colorectal tumorigenesis, implicating LRRK2 as a potential target in colon cancer patients with hyper LRRK2 kinase activity.

## Introduction

Leucine-rich repeat kinase 2 (LRRK2) is a serine-threonine protein kinase with multiple domains and belongs to the ROCO protein family. It consists of an N-terminal armadillo domain, Ankyrin repeats, a leucine-rich repeats (LRR) domain, a Ras of complex protein domain (Roc) GTPase, a C-terminal of Roc (COR) domain, a kinase and a C-terminal WD40 domain (1,2). LRRK2 plays numerous physiological roles in autophagy, lysosome function, endocytosis, and Golgi network modulation (3-5). Emerging evidence has further suggested that LRRK2 is an important regulator of immune responses (6,7). We recently identified LRRK2 as a novel modulator of the activation of NLRC4 inflammasome (8). Of note, LRRK2 was first identified in 2004 as a genetic cause of Parkinson’s disease (PD) (9,10). Subsequently, LRRK2 mutations were also discovered to be linked with inflammatory bowel disease (IBD) (11-13), cancer, and leprosy et al (14,15). Among these, LRRK2 G2019S is the most prevalent variant associated with PD. It is found in a wide range of ethnic groups, in 1-3% of sporadic and 4-8% of familial PD cases (16,17). Intriguingly, several previous studies have demonstrated that LRRK2 G2019S mutation carriers have an overall elevated risk of cancer, especially hormone-related cancers, and breast cancer in women (18-20). Recent studies further suggested that LRRK2 G2019S PD patients exhibit drastically increased risks of colon cancer and leukemia compared with idiopathic PD patients (21,22). However, the underlying mechanisms of the genetic susceptibility of LRRK2-G2019S to tumorigenesis remain largely unknown. There is, therefore, a critical need to experimentally test whether and how this mutation promotes cancer development in the colon.

Colorectal cancer is the third most prevalent cancer worldwide (21). Inflammation is recognized as a hallmark of cancer development and progression (23). In humans, the correlation between IBD and colorectal cancer has long been established. Colitis-associated colorectal cancer (CAC) may reach as many as 18.4% at 30 years after the onset of ulcerative colitis (24), which is 2 to 3-fold that of the general population (25). Furthermore, CAC patients have a worse prognosis than colorectal cancer patients without a history of IBD (26,27). Increased mortality rates for CAC have also been reported (28,29). While molecular mechanisms underlying colorectal cancer remain unclear, inflammation is considered a driving force for the pathogenesis (30). Azoxymethane/dextran sulfate sodium (AOM/DSS)-induced colon cancer in mice is a widely used animal model for investigating the pathophysiology of CAC.

LRRK2 is expressed in myeloid cells, B cells, T cells, microglial cells, and epithelial cells (31,32). LRRK2 is upregulated in the lamina propria of intestinal biopsies from patients with Crohn’s disease (CD) (33). GWAS has demonstrated the association between LRRK2 locus and IBD (34,35). Further exome sequencing uncovered the presence of shared LRRK2 alleles in CD and PD, such as gain-of-function variant N2081, which is located in the kinase domain of LRRK2 (36). LRRK2 G2019S is a gain-of-kinase activity mutant as well. We thus hypothesized that LRRK2 G2019S promotes the development of colorectal cancer by promoting intestinal inflammation.

In the present study, we report that LRRK2 G2019S promotes colon tumorigenesis in a mouse model of CAC. Furthermore, we revealed that LRRK2 G2019S increases mouse susceptibility to DSS-induced colitis. Mechanistically, LRRK2 G201S drastically enhances inflammasome activation, triggers epithelial cell necrosis, and induces the production of reactive oxygen species in the gut, and the kinase activity of LRRK2 is important for these functions. Therefore, we have experimentally demonstrated the pro-tumorigenic role of LRRK2 G2019S in the pathogenesis of colorectal cancer.

## Materials and Methods

### Animals

LRRK2 G2019S KI (LRRK2 KI) mice (37) were all purchased from Jackson Laboratory (JAX030961). To get littermate wild type control mice, LRRK2^KI/+^ male mice were mated with LRRK2^KI/+^ female mice, sex-matched LRRK2^+/+^ (LRRK2 WT) and LRRK2^KI/KI^ (LRRK2 KI) mice were used at the age of 8-12 weeks. Both male and female mice were used for the experiments. All mice were bred and maintained in individually ventilated cages under specific pathogen-free conditions in accredited animal facilities. Animal experiments were approved by the Institutional Animal Care and Use Committee of the University of Iowa.

### DSS-induced acute colitis

Colitis was induced by treating mice with 2.5% DSS for 7 days as described previously (38). In brief, sex-matched LRRK2 WT and LRRK2 G2019S KI mice at the age of 8–12 weeks were given 2.5% DSS (MP Biomedical) in normal drinking water for 7 days, followed with regular drinking water from animal facility for another 2 days to induce acute colitis. Mouse weights were monitored every day during the colitis model. The humane endpoints for DSS study, in accordance with Institutional Animal Care and Use Committee recommendations from the University of Iowa, was the loss of 20% of initial body weight.

### AOM/DSS-induced colon cancer

Colonic tumors were induced using the azoxymethane/dextran sulfate sodium (AOM/DSS) protocol as described previously (39,40). Male and female mice aged 8-12 weeks were utilized for the study. The mice were initially injected intraperitoneally with AOM at a dose of 10 mg/kg body weight. On day 3 after AOM administration, the mice were exposed to 2.5% DSS in their drinking water for 5 consecutive days. This was followed by a resting period of 16 days. This cycle was repeated for another two times. Tumorigenesis assessment was performed on day 65 following the initial AOM injection. For tumor analysis, after mouse euthanasia, the colon was longitudinally cut and then fixed in 10% neutral buffered formalin overnight. All colon tumors were carefully counted and measured using a stereo microscope. Representative tumors were selected for paraffin embedding and sectioning at a thickness of 5 μm. Histological analyses were conducted by hematoxylin and eosin (H&E) staining.

### LRRK2-IN-1 administration

The LRRK2 kinase inhibitor LRRK2-IN-1 (MedChemExpress) was reconstituted in corn oil as described previously (41). Experimental groups of mice were treated with LRRK2-IN-1 by intraperitoneal (IP) injection daily at a dose of 100 mg/kg body weight and for a duration of 7 days. This inhibitor treatment was administered concurrently with the DSS treatment.

### Immunoblot

Tissues were homogenized using radioimmunoprecipitation (RIPA) assay buffer, which was supplemented with Complete Mini Protease Inhibitor Cocktail and Phosphatase Inhibitor Cocktail from Roche. The lysates were put on ice for a period of 30 minutes and vortexed every 5 minutes. Following centrifugation at 15,000 rpm for 15 minutes at 4°C, the supernatants were collected. Protein concentration was determined by a BCA Protein Assay Kit from Pierce. Subsequently, the proteins were resolved by SDS-PAGE and transferred to a 0.45-mm PVDF membrane. For immunoblot analysis, the indicated primary antibodies are listed in supplemental Table 1 and were used at 1000-fold dilution. Horseradish peroxidase (HRP)-conjugated secondary antibodies were used depending on the host species of the primary antibodies.

### ELISA

Whole colon cultures derived from DSS-treated mice were used to assess cytokine production using ELISA kits obtained from R&D Systems, following the manufacturer’s instructions. The following catalog numbers were used for specific cytokines: TNF (DY410-05), IL-1β (DY401-050), IL-6 (DY406-05), and IL-18 (7625-05). Cytokine levels were normalized to the weight of the colon tissue used.

### Intestinal permeability analysis

Intestinal permeability in vivo was assessed using a FITC-dextran (average molecular mass 4000 kDa; Sigma-Aldrich) gavage experiment, both in naive conditions and during colitis. Briefly, mice were fasted overnight prior to gavage of FITC-dextran at a dosage of 60 mg per 100 g of body weight. After 4 hours, blood samples were collected via cardiac puncture and sera were obtained by centrifugation at 10,000 × g for 10 minutes. The fluorescence intensity of the serum samples was measured using a SpectraMax i3 instrument from Molecular Devices. The concentration of FITC-dextran was determined by referencing a standard curve generated from serial dilutions of FITC-dextran.

### Isolation of Intestinal epithelial cells (IECs)

IECs were isolated as described previously (42). The procedure involved gently extracting the large intestine from the abdominal cavity, followed by the removal of mesentery and fatty tissue using forceps. The large intestine, excluding the cecum, was dissected, and longitudinally opened, then cut into 1 cm pieces. The intestinal pieces were thoroughly washed three times with ice-cold PBS. Subsequently, the pieces were further chopped into 5 mm fragments and incubated in 5 mM EDTA/PBS solution at 4°C on a rocking platform for 30 minutes. IECs were released by shaking the tubes for 2 minutes, and then collected by centrifugation at 200 × g for 10 minutes at 4°C. The dissociated cells were washed and resuspended in PBS with 10% FBS.

### Total ROS and Mitochondrial ROS measurement

To assess the total ROS in IECs, the cells were incubated with 10 mM CM-H2DCFDA (Life Technologies, # C6827) which is a cell-permeable indicator for ROS. For the detection of mitochondrial ROS levels, the isolated IECs were incubated with Mitosox (M36008, Life Technologies) for 15 minutes following the manufacturer’s instructions. Subsequently, the fluorescence levels were quantified by flow cytometry. All flow data were analyzed by FlowJo software.

### Colonic explants

Whole colons were harvested from LRRK2 WT or LRRK2 G2019S KI mice, thoroughly rinsed with serum-free DMEM medium, and weighed to determine their initial weight. The collected colon tissues were cut into 2 mm pieces and then cultured as explants in regular RPMI 1640 medium supplemented with 10% FBS, L-glutamine, penicillin, and streptomycin, and placed in a standard cell-culture incubator for 24 hours. After the culture, the cell-free supernatants were obtained by centrifuging at 12,000 × g for 10 minutes at 4°C and stored in aliquots at −20°C for further analysis.

### Quantitative PCR

Whole colon tissues or cells were carefully preserved and homogenized using TRIzol reagent (Invitrogen) to ensure optimal RNA extraction. The RNA extraction procedure was carried out following the manufacturer’s instructions, and the extracted RNA was promptly reverse-transcribed into complementary DNA (cDNA). Quantitative PCR (Q-PCR) analysis was performed using SYBR Green Real-time PCR Master Mix on a Real-Time PCR System (Applied Biosystems). The primer sequences utilized for the Q-PCR amplification are provided in the supplemental Table 2.

### Immunohistochemistry

Formalin-fixed and paraffin-embedded colon sections or tumor samples were meticulously deparaffinized and rehydrated. Antigen retrieval was performed using citrate antigen retrieval buffer, followed by gradual cooling to room temperature. Following permeabilization and blocking steps, the sections were incubated overnight at 4°C with primary antibodies, including anti-cyclin D1 (dilution, 1:200, CST), anti-p-STAT3 (dilution, 1:200, CST), and anti-Ki67 (dilution, 1:500, CST). Subsequently, the sections were incubated with fluorescence-conjugated secondary antibodies. The images were taken by fluorescent microscope (Olympus, model DP74-CU).

### Histological analysis

For H&E staining, approximately 3 mm sections of colon tissues were meticulously fixed in 10% formalin or 4% paraformaldehyde and subsequently embedded in paraffin. The paraffin-embedded sections were then stained with H&E to facilitate histological analysis. Scoring of the sections was performed in a scale ranging from 0 to 10 by summarizing the scores of the severity of inflammation, the extent of inflammation, and crypt damage as adapted from previous study (43). In brief, the severity of inflammation was scored on a scale of 0-3, with 0 indicating no inflammation, 1 denoting mild inflammation, 2 representing moderate inflammation, and 3 indicating severe inflammation. The extent of inflammation was also scored on a scale of 0-3, with 0 indicating no inflammation, 1 denoting involvement of the mucosa, 2 representing involvement of both the mucosa and submucosa, and 3 indicating transmural/muscularis/serosa involvement. Crypt damage was scored on a scale of 0-4, with 0 indicating no damage, 1 representing one-third crypt damage, 2 denoting two-thirds crypt damage, 3 indicating loss of crypts with surface and epithelium still present, and 4 denoting complete loss of crypts and surface epithelium.

### Subcellular fractionation protocol

Cytosol and mitochondria fragments were isolated using a Mitochondria/Cytosol Fractionation Kit (ab65320). In brief, cells were treated with 500 μL of fractionation buffer and incubated on ice for 15 minutes. To ensure cell lysis, the cell suspension was passed through a 27-gauge needle 10 times using a 1 mL syringe or until all cells were lysed. The lysed sample was then left on ice for an additional 20 minutes. The lysed sample was centrifuged at 3,000 rpm for 5 minutes; the resulting pellet contained nuclei while the supernatant contained cytoplasm, membranes, and mitochondria. Then the supernatant was carefully transferred into a fresh tube and subjected to centrifugation at 8,000 rpm for 5 minutes, then the resulting pellet contained mitochondria. The supernatant, which contained the cytoplasm and membrane fraction, was transferred into a fresh tube, and stored for further analysis. The subcellular fractionation buffer used in the protocol included 20 mM HEPES (pH 7.4), 10 mM KCl, 2 mM MgCl_2_, 1 mM EDTA and 1 mM EGTA. Additionally, 1 mM DTT was added, and 1 piece of PI cocktail was added per 10 mL of buffer just before use.

### Statistic

The p values of weight loss comparison were determined by two-way ANOVA as specified in the figure legend. p values for two group comparisons were determined by Student’s t test. Unless otherwise specified, all results are shown as mean ± SD. p value<0.05 was considered significant.

## Results

### LRRK2 G2019S promotes the pathogenesis of colitis-associated cancer

Previous studies have provided evidence that individuals carrying the LRRK2 G2019S mutation were at increased risk of developing cancers (21,22). However, the direct impact of this mutation on cancer development has not been tested to date. Here, we utilized a mouse model of CAC (39,40) to test the role of LRRK2 G2019S in colon tumorigenesis in LRRK2 G2019S KI (LRRK2 KI) mice. In this model, mice receive a single treatment of AOM by intraperitoneal injection, followed by three cycles of DSS treatment in regular drinking water (**Fig. 1A**). After AOM/DSS induction, both LRRK2 KI mice and WT controls developed colon tumors and the tumors were exclusively clustered in the distal colon (**Fig. 1B**). Some of the LRRK2 KI mice exhibited severe rectal bleeding, diarrhea, or weight loss toward the end of the treatment. Additionally, the average number of tumors per mouse in LRRK2 KI mice (20 ± 3.85) was about double of that in WT counterparts (9.8 ± 2.56) (**Fig. 1C**). The presence of LRRK2 G2019S also influenced the size of the tumors. Approximately 30.6% of the polyps developed in WT mice were small adenomas (2-5 mm diameter), in contrast with 45% in LRRK2 KI mice. Furthermore, 6.1% in WT mice versus 11% polyps in LRRK2 KI mice developed into large tumors (>5 mm diameter) (**Fig. 1D and Fig. 1E**). Similarly, the average tumor load, calculated by summing the diameters of all tumors in each mouse, was significantly higher in LRRK2 KI mice compared to WT mice (**Fig. 1F**). Further histological analyses revealed that the majority lesions formed in WT mice were either polypoid colonic tissue or tubular adenomas and most with low-grade dysplasia, featuring basally located, elongated, hyperchromatic nuclei. There was no lamina propria invasion. In contrast, the large tumors formed in LRRK2 KI mice exhibited extensive and confluent high-grade dysplasia, loss of nuclei polarity, and lamina propria invasion, characteristic of adenocarcinoma (**Fig. 1G**). Tumorigenesis can be divided into three mechanistic stages: initiation (genomic alteration), promotion (proliferation of genetically altered cells), and progression (tumor growth) (44). Therefore, these data indicate that LRRK2 G2019S not only enhances tumor promotion but also increases tumor progression.

**Figure 1.**
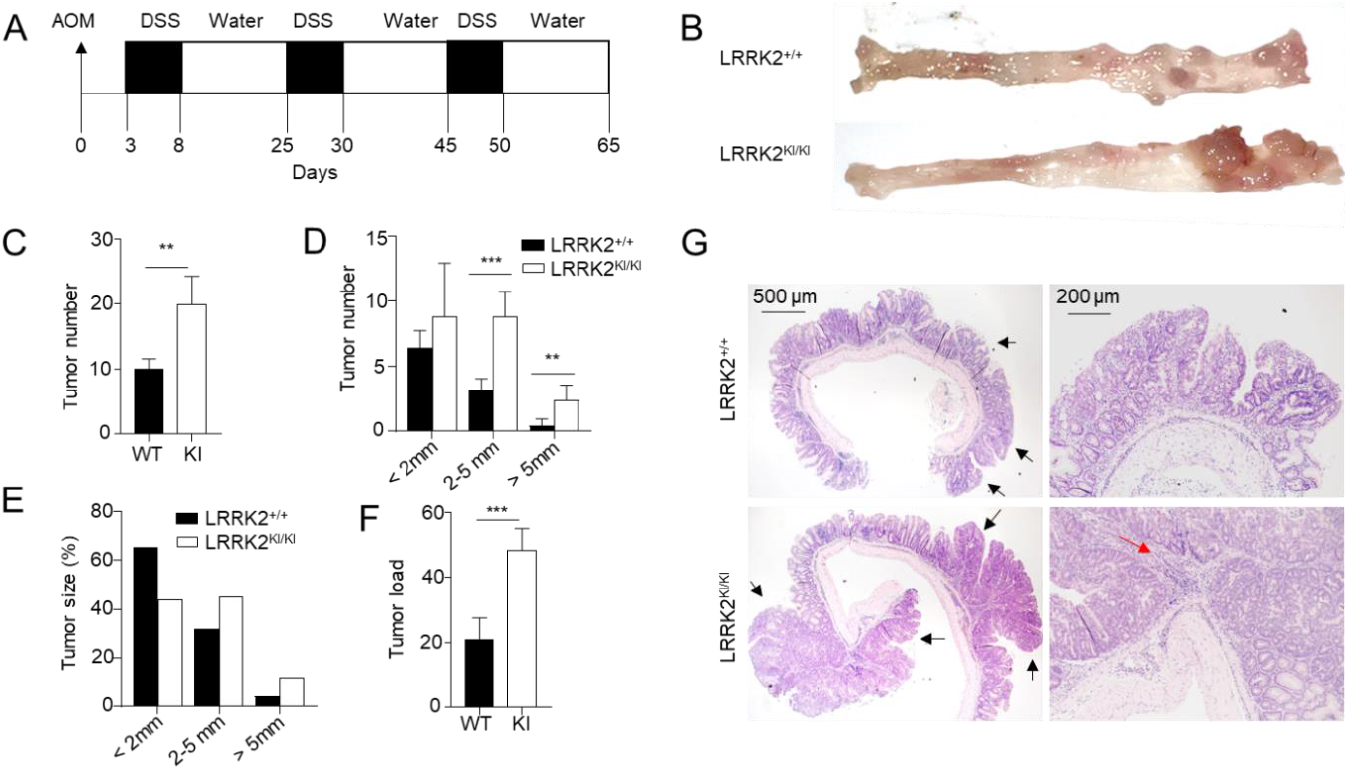
LRRK2 G2019S promotes the pathogenesis of colitis-associated colon cancer. (A) Schematic diagram for AOM/DSS-induced inflammatory carcinogenesis. 8-12 weeks-old mice were injected with AOM at a dose of 10 mg/kg mouse by intraperitoneal (IP) injection. 3 Days later, mice were given 2.5% DSS for 5 days, then mice were rested for 16 days. This process was repeated for another two cycles starting on day 25 and day 45, respectively. Mice were euthanized on day 65 for analysis as stated below. (B) Representative image of gross colons from LRRK2^+/+^ and LRRK2^KI/KI^ mice. (C) Tumor numbers in colons of LRRK2^+/+^ (WT) and LRRK2^KI/KI^ (KI) mice. (D) Tumor size distributions in colons of LRRK2^+/+^ and LRRK2^KI/KI^ mice. (E) Tumor size percentage in colons of LRRK2^+/+^ and LRRK2^KI/KI^ mice. (F) Average tumor load was determined by summing all tumor diameters for a given animal. (G) H&E staining of tumor sections from colons of LRRK2^+/+^ and LRRK2^KI/KI^ mice. Black arrows point to the tumors. The red arrow points to tumor cells invading lamina propria. p values were determined by Student’s t test, n=5 mice/group. Data represent mean ± SD. *p<0.05, **p<0.01 and ***p<0.001. Data are representative of three independent experiments.

### LRRK2 G2019S promotes inflammation and cell proliferation in tumors

To elucidate the molecular mechanisms underlying the impact of LRRK2 G2019S on CAC development, we examined colon tumor tissues from AOM/DSS-treated LRRK2 KI and WT mice to assess the activation of various pro-tumor effectors. First, we compared the pro-inflammatory gene expression in the colon tumor tissues using real-time PCR. The result revealed much higher expression of inflammatory genes including IL-1β, IL-6, IL-11, IL-17, IL-23 and COX-2 et al; while expression of TNF, CCL7 and CXCL9 were comparable between the two groups (**Fig. 2A)**. Moreover, factors associated with tissue remodeling and angiogenesis, such as Ang4, VEGF, Wnt5a and MMP10, were also significantly upregulated in the LRRK2 KI mice (**Fig. 2B**).

**Figure 2.**
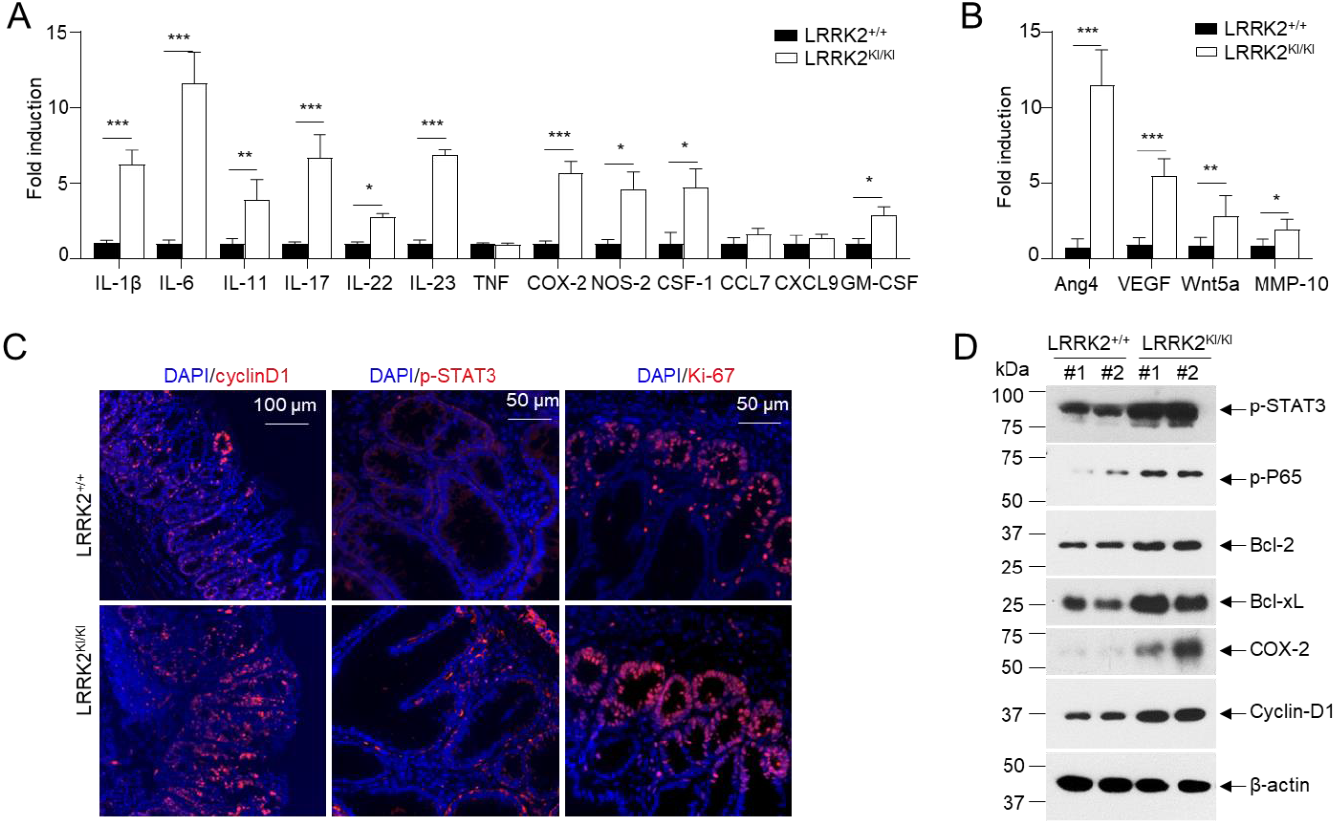
LRRK2 G2019S promotes inflammation and cell proliferation in colon tumors. 65 days after AOM/DSS induction, LRRK2^+/+^ and LRRK2^KI/KI^ mice were euthanized, and colon tumor tissues were collected for the following analysis: (A) Real-time PCR analysis of Inflammatory genes in colon tumors. (B) Real-time PCR analysis of pro-tumorigenic genes in colon tumors. (C) Fluorescence staining of tumor tissues with indicated antibodies and counterstained by DAPI. (D) Immunoblot analysis of key proteins involved in colon tumorigenesis as indicated. Lysates were prepared from colon tumor tissues of LRRK2^+/+^ and LRRK2^KI/KI^ mice. Numbers represent individual mouse from each group. p values were determined by Student’s t test, n=5 mice/group. Data represent mean ± SD. *p<0.05, **p<0.01 and ***p<0.001. Data are representative of three independent experiments.

IL-1β has been shown to play an important role in CAC development via activating NF-κB (45). IL-6, a well-established cytokine induced by IL-1β in intestinal epithelial cells (IECs) (46), has been shown to promote tumor progression in CAC through the activation of the oncogene STAT3 (47) in addition to IL-11 (48). Consistent with these, we did observe increased staining of phosphorylated STAT3 in the colon IECs of LRRK2 KI mice (**Fig. 2C**). The activation of STAT3 or NF-κB upon AOM/DSS induction were expected to promote proliferation and survival of IECs, then we tested the proliferation markers in the colon epithelium of LRRK2 KI and WT mice. Cyclin D1, which is a target gene of both STAT3 and NF-κB and critical for cell proliferation and survival, was highly expressed in the epithelium of LRRK2 KI mice compared to WT mice; Ki-67 protein has been widely used as a proliferation marker for tumor cells. Consistently, we detected a higher number of Ki-67-positive cells in the colon epithelium of LRRK2 KI mice compared to WT controls (**Fig. 2C**). Consistent with gene expression analysis and immunohistochemical staining, immunoblot analysis revealed p-STAT3, p-P65, Bcl-2, Bcl-xL and Cyclin-D1 levels in tumor tissues from KI mice were dramatically increased. Cox-2 is important to produce prostaglandin E2 (PGE2) during inflammation. Of note, PGE2, which is the most abundant prostanoid in colorectal cancer, promotes anti-tumor immune responses by inducing Treg and MDSCs (30) and promotes tumor initiation and growth by upregulating the expression of DNMT1 and DNMT3B (49). The immunoblot assay suggested increased level of COX-2 in the tumor tissues from LRRK2 KI mice (**Fig. 2D**). Taken together, these results demonstrate that the activation of IL-1β-NF-κB and IL-6/IL-11-STAT3 pathways may potentially play pivotal roles in the promotion of tumorigenesis in the colons of LRRK2 KI mice.

### LRRK2 G2019S KI mice are highly susceptible to DSS-induced colitis

The association between chronic inflammation and the development of colorectal cancer has been extensively documented (50,51). To unravel the mechanisms underlying the increased tumor promotion and progression observed in LRRK2 KI mice, we hypothesized that this mutation could enhance susceptibility to DSS-induced colitis, thus promoting cancer pathogenesis. To test this, we utilized the DSS-induced colitis model to investigate the susceptibility of LRRK2 KI mice to intestinal inflammation. Both LRRK2 KI mice and WT control mice were subjected to 2.5% DSS treatment in drinking water for 7 consecutive days. Notably, LRRK2 KI mice exhibited more rapid weight loss (**Fig. 3A**). In addition, LRRK2 KI mice manifested more exacerbated colitis symptoms including shortened colon length and increased spleen weight compared with those in WT mice (**Fig. 3B**). Analysis of supernatants from colonic explant culture revealed much higher levels of the cytokines of IL-6 and TNF-α in LRRK2 KI mice compared with WT controls (**Fig. 3C**). Additionally, histology analysis found that LRRK2 KI mice displayed heightened gut damage and inflammatory cell infiltration following DSS treatment compared with WT controls (**Fig. 3D-E**). The enhanced inflammatory response in LRRK2 KI mice was associated with exacerbated impairment of intestinal epithelial integrity, as demonstrated by the increased leakage of FITC-dextran from the gastrointestinal tract into the systemic circulation (**Fig. 3F**). Consistent with the observed tissue damage and inflammation, we also identified highly upregulated expression of pro-inflammatory cytokines and chemokines, such as IL-1β, IL-6, IL-11, IL-17, CXCL9 and COX-2 et al, in the colon of LRRK2 KI mice compared with WT controls (**Fig. 3G**). However, the expression of pro-tumorigenic genes including Ang4, VEGF, Wnt5a and MMP-10 were comparable between LRRK2 KI and WT control groups (**Fig. 3H**), suggesting that LRRK2 G2019S may promote tumor development primarily by promoting inflammation. Notably, we didn’t detect any difference in colon structure by H&E staining and pro-inflammatory cytokines expression detected by Q-PCR when the LRRK2 KI and WT controls were at homeostatic status without any treatment (data not shown), suggesting LRRK2 G2019S doesn’t affect the baseline of gut inflammation. Taken together, our findings indicate that LRRK2 KI mice are more susceptible to DSS-induced colitis than WT counterparts, and LRRK2 G2019S may promote tumorigenesis through promotion of inflammation.

**Figure 3.**
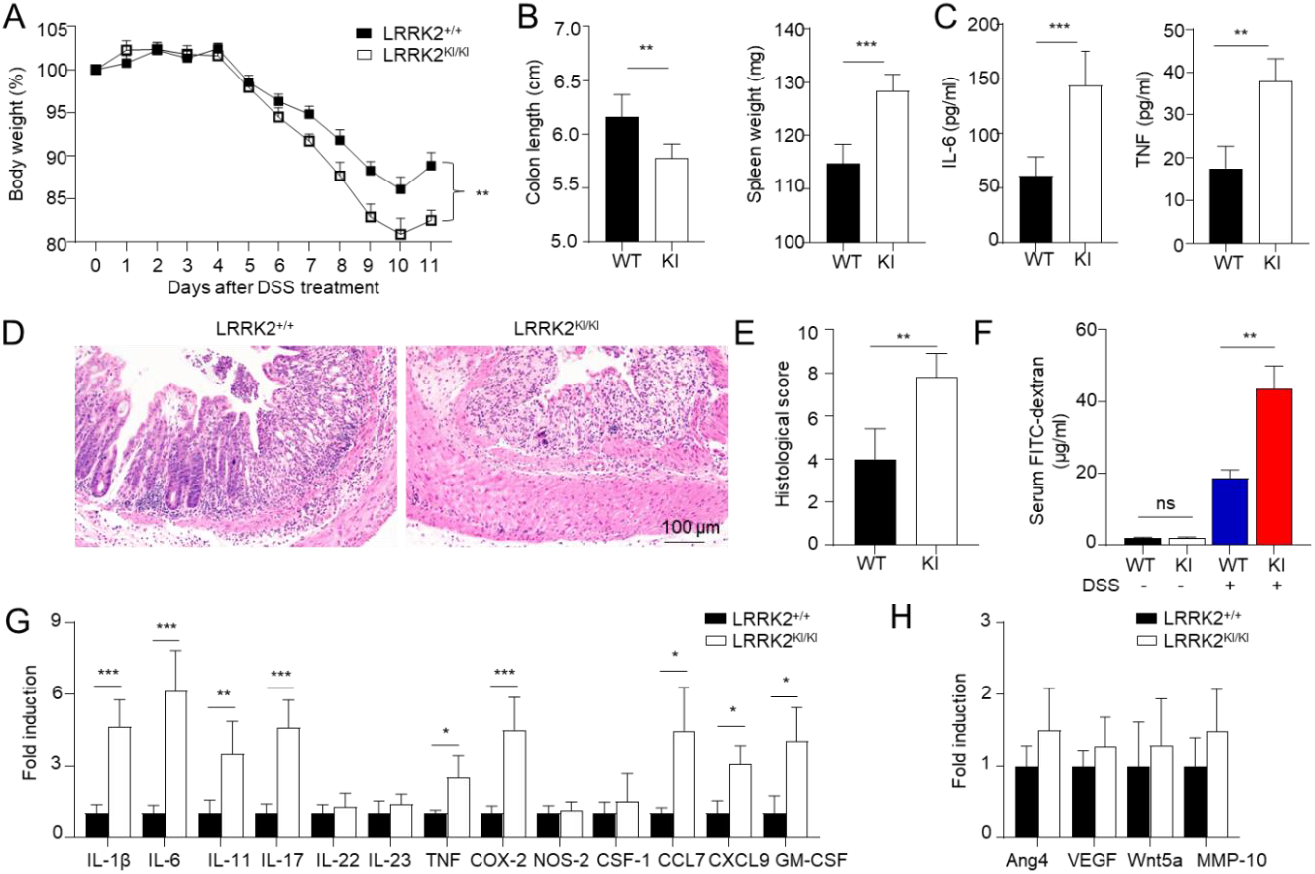
LRRK2 G2019S KI mice are highly susceptible to DSS-induced colitis. Acute colitis was induced in LRRK2^+/+^ (WT) and LRRK2^KI/KI^ (KI) mice with 2.5% DSS in drinking water for 7 days. Mice were euthanized on day 10. Colitis severity was shown by: (A) Body weight loss; (B) Colon length and spleen weight; (C) IL-6 and TNF levels in colon explant culture (100 mg colon tissue/ml medium) analyzed by ELISA; (D-E) H&E staining of colonic tissue and histology score; (F) Gut permeability assay. Untreated or DSS-treated LRRK2^+/+^ and LRRK2 ^KI/KI^ mice were gavaged by FITC-dextran, 4h later sera were collected, and FITC-dextran level was measured. (G) Real-time PCR analysis of Inflammatory genes in the colon tissues. (H) Real-time PCR analysis of pro-tumorigenic genes in colon tissues. n=5/group except panel A (n=10/group). P values were determined by two-way ANOVA in panel A and student’s t test in other related panels. Data represent mean ± SD. *p<0.05, **p<0.01 and ***p<0.001. Data are representative of three independent experiments.

### Kinase activity of LRRK2 G2019S is critical for the exacerbated colitis

Previous studies have demonstrated the suppressive effects of the LRRK2 kinase inhibitor LRRK2-IN-1 on cytokine production in vitro and colitis in vivo (41). To examine the potential role of LRRK2 kinase activity in DSS-induced colitis, we then investigated the impact of LRRK2-IN-1 treatment on colitis development in both LRRK2 KI mice and littermate controls using the DSS colitis model. Remarkably, administration of LRRK2-IN-1 inhibited DSS-induced colitis in both LRRK2 KI and WT groups (**Fig. 4**). First, we found the body weight loss induced by DSS was drastically prevented by the inhibitor not only in LRRK2 KI groups but also in the WT groups (**Fig. 4A**). Treatment with LRRK2-IN-1 alleviated the reduction in colon length, which is indicative of the severity of colon inflammation. Enlarged spleen weight, indicating a system inflammation induced by DSS, was also relieved by this potent inhibitor (**Fig. 4B**), suggesting LRRK2-IN-1 played an important role in dampening the intestinal inflammation. Moreover, histology analysis revealed that LRRK2-IN-1 treatment alleviated colon damage (**Fig. 4C** and **Fig. 4D**). Furthermore, inhibition of LRRK2 kinase activity partially mitigated colonic inflammation at molecular level detected by Q-PCR in both LRRK2 KI and WT groups (**Fig. 4E**). In total, these findings suggest that LRRK2 kinase activity plays a crucial role in the development of DSS-induced colitis, targeting LRRK2 kinase activity might provide a potential method for inhibiting the development of colitis and colitis-associated cancer.

**Figure 4.**
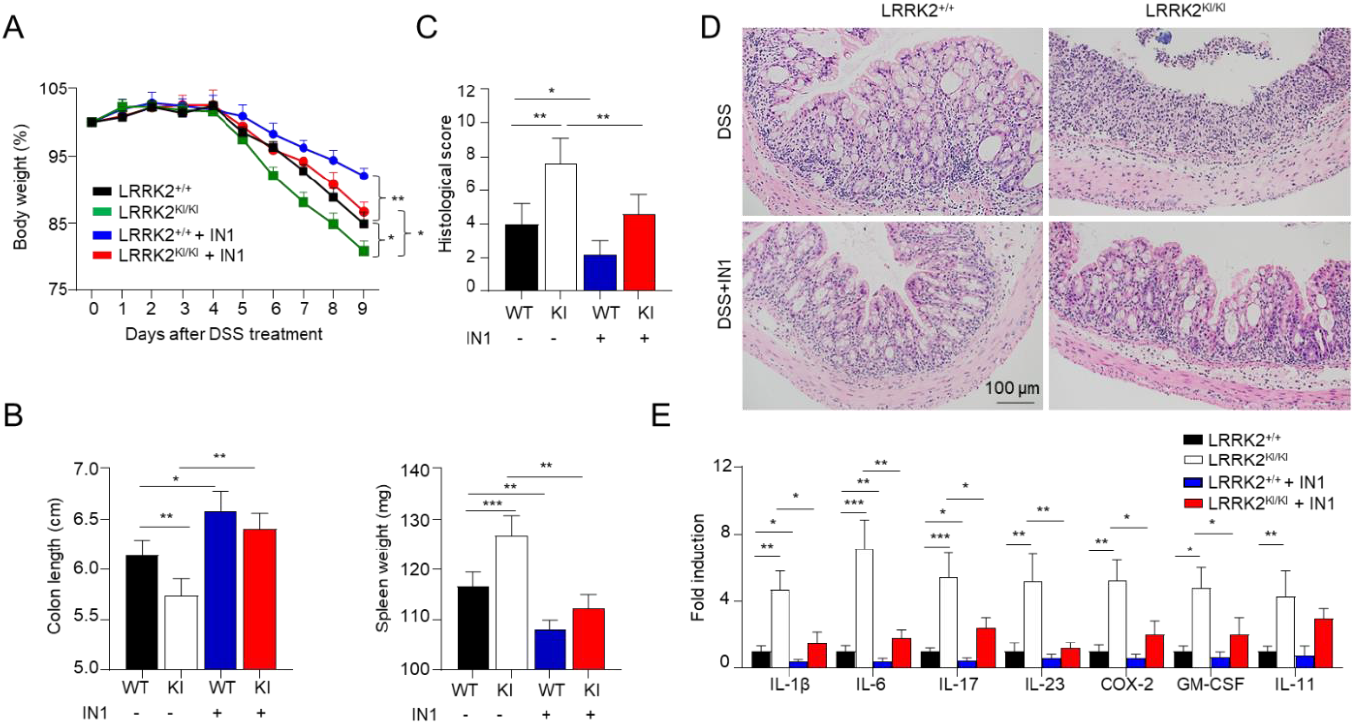
Kinase activity of LRRK2 G2019S is critical for the exacerbated colitis. Acute colitis was induced in LRRK2^+/+^ (WT) and LRRK2^KI/KI^ (KI) mice with 2.5% DSS in drinking water. LRRK2-IN1(IN-1) inhibitors were used to treat both the WT and KI mice at a dose of 100 mg/kg every day by IP injection, mice treated with vehicles were included as controls as indicated. Mice were euthanized on day 9. Colitis severity was shown by: (A) Body weight loss; (B) Colon length and spleen weight; (C-D) H&E staining of colonic tissue and histology score; (E) Real-time PCR analysis of Inflammatory gene expression as indicated in colon tissues. n=5 mice/group. P values were determined by two-way ANOVA in panel A and Student’s t-test in other related panels. Data represent mean ± SD. *p<0.05, **p<0.01 and ***p<0.001. Data are representative of three independent experiments.

### LRRK2 G2019S promotes inflammasome activation and necrosis in the gut epithelium

Inflammasome activation plays an important role in gut homeostasis and IBD (52). We have recently demonstrated that LRRK2 is an upstream regulator of NLRC4 inflammasome activation (8). Of note, gain-of-function mutations of NLRC4 result in hyperactivation of the inflammasome and lead to autoinflammation and enterocolitis (53,54). These findings implicate that LRRK2 G2019S may play an important role in inflammasome activation in the DSS colitis model. Therefore, we determined the status of inflammasome activation in colon tissues from LRRK2 KI mice and WT controls after colitis induction. General markers for inflammasome activation include the production of mature/secreted IL-β and IL-18, caspase-1 cleavage, and Gasdermin D (GSDMD) cleavage which leads to cell pyroptosis. First, we measured the secretion of IL-β and IL-18 in the supernatants of colonic explant culture from LRRK2 KI and WT controls by ELISA, both IL-1β and IL-18 levels were much higher in supernatants from LRRK2 KI mice compared with those from WT controls (**Fig. 5A**). Second, we utilized immunoblot to analyze the inflammasome activation in the gut epithelium by isolating IECs from colons of LRRK2 KI mice and WT controls, we observed increased cleavage of IL-1β, caspase-1 and GSDMD in the IECs from LRRK2 KI mice compared with WT counterparts (**Fig. 5B**), suggesting more robust inflammasome activation in the gut epithelium of LRRK2 KI mice after DSS treatment.

**Figure 5.**
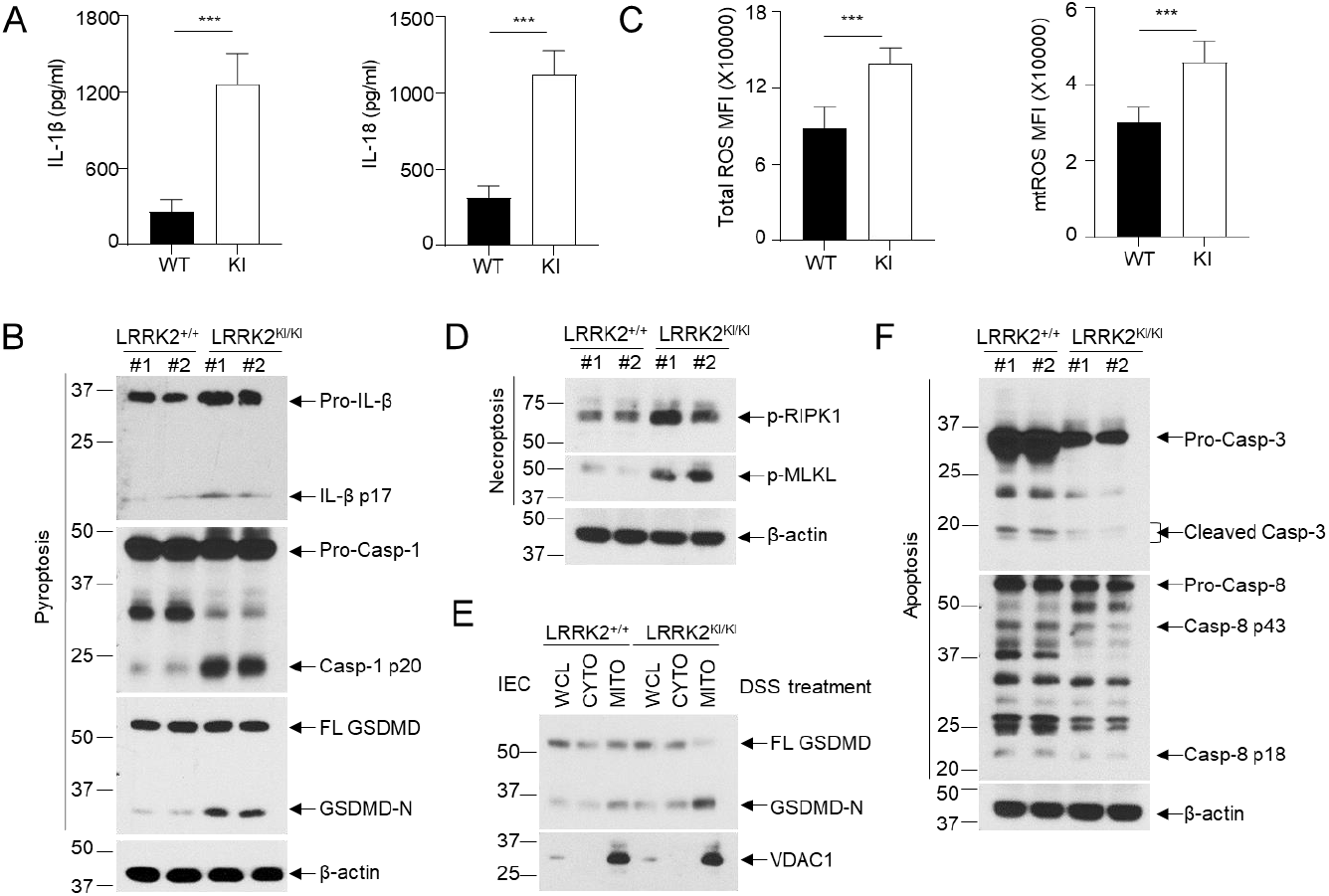
LRRK2 G2019S promotes inflammasome activation and necrosis in the gut epithelium. (A) IL-1β and IL-18 in colon explant culture of LRRK2^+/+^ (WT) and LRRK2^KI/KI^ (KI) mice were measured by ELISA. IECs from LRRK2^+/+^ and LRRK2^KI/KI^ mice were isolated 9 days after colitis induction and were collected for the following experiments: (B) Inflammasome activation in IECs was analyzed by immunoblot as indicated. (C) Mean fluorescence intensities (MFI) of total ROS and mitochondrial ROS (mtROS) from IECs are shown as indicated. ROS levels were analyzed by flow cytometry. (D) Necroptosis in IECs were analyzed by immunoblot as indicated. (E) GSDMD-N levels in cellular fractions of IECs were analyzed by immunoblot as indicated. IECs were lysed, then whole cell lysates (WCL) were fractionated into cytoplasm (CYTO) and mitochondria (MITO). (F) Apoptosis in IECs was analyzed by immunoblot as indicated. p values were determined by Student’s t test. n=5 mice/group. Data represent mean ± SD. ***p<0.001. Data are representative of two independent experiments.

ROS has been found to be increased in IBD patients, it is also a well-established factor in promoting inflammasome activation (55). A recent study suggested that LRRK2 G2019S promotes mitochondrial ROS (mtROS) production in bone marrow-derived macrophages (BMDMs) with transgenic LRRK2 G2019S expression (56). Furthermore, the increased mtROS mobilizes cleaved GSDMD-N to mitochondrial and form pores in the membrane which further enhance the release of mtROS. The excessive ROS in LRRK2 G2019S transgenic BMDMs promotes necroptosis instead of pyroptosis (56). We then wondered whether LRRK2 G2019S promotes ROS production and necroptosis in the gut epithelium. Intriguingly, our data indicated an increase in both total ROS and mitochondrial ROS levels in IECs from LRRK2 KI mice compared with WT controls after DSS treatment (**Fig. 5C**). With that, we further tested the necroptosis of IECs from LRRK2 KI mice and WT controls by immunoblot analysis. We found a marked increase of phosphorylated-RIPK1 and MLKL (**Fig. 5D**), suggesting elevated necroptosis in the gut epithelium of LRRK2 KI mice. Interestingly, we also observed the increased translocation of GSDMD to mitochondria of the IECs in LRRK2 KI mice following DSS treatment (**Fig. 5E**), suggesting that G2019S promotes GSDMD-N location in mitochondrial may be universal and not limited to a specific cell type. In contrast to necroptosis, we didn’t observe the increase of apoptosis, characterized by caspase 3 and caspase 8 activation, in the gut epithelium of LRRK2 KI mice after DSS treatment (**Fig. 5F**). Taken together, our data demonstrate that LRRK2 G2019S KI promotes inflammasome activation and necrosis in the context of DSS-induced colitis. These findings shed light on the underlying mechanisms through which LRRK2 G2019S may contribute to the pathogenesis of colon cancer.

### LRRK2 G2019S promotes inflammation and cell proliferation during the early stage of tumorigenesis

To gain further insights into how LRRK2 G2019S contributes to increased tumor promotion and progression, we conducted a comparative analysis of the inflammation and proliferation status in the colon tissues of LRRK2 KI mice and WT control mice during the early stage of tumorigenesis following AOM and DSS treatment (**Fig. 6A**). After AOM administration and the first cycle of DSS treatment, we found a much stronger induction of the inflammatory genes of IL-1β, IL-6, and COX-2 in colon tissues of LRRK2 KI mice compared to those in WT controls on day 8 and day 15 after AOM+DSS treatment, but these genes were not primed by AOM alone three days after its administration (**Fig. 6B**). In contrast, the pro-tumorigenic genes of Ang4 and VEGF were found to be more robustly expressed in the colon of LRRK2 KI mice three days after AOM treatment, and further increased in both groups after DSS treatment but more dramatically upregulated in LRRK2 KI mice (**Fig. 6C**). Consistent with the enhanced expression of proinflammatory and pro-tumorigenic genes in the colon tissue of LRRK2 KI mice, immunofluorescent staining uncovered an increased number of Ki-67-positive cells in the colon epithelium of LRRK2 KI mice 15 days after AOM/DSS administration (**Fig. 6D**), indicating elevated cell proliferation of IECs in the gut of LRRK2 KI mice during the early stages of AOM/DSS induction compared with WT controls.

**Figure 6.**
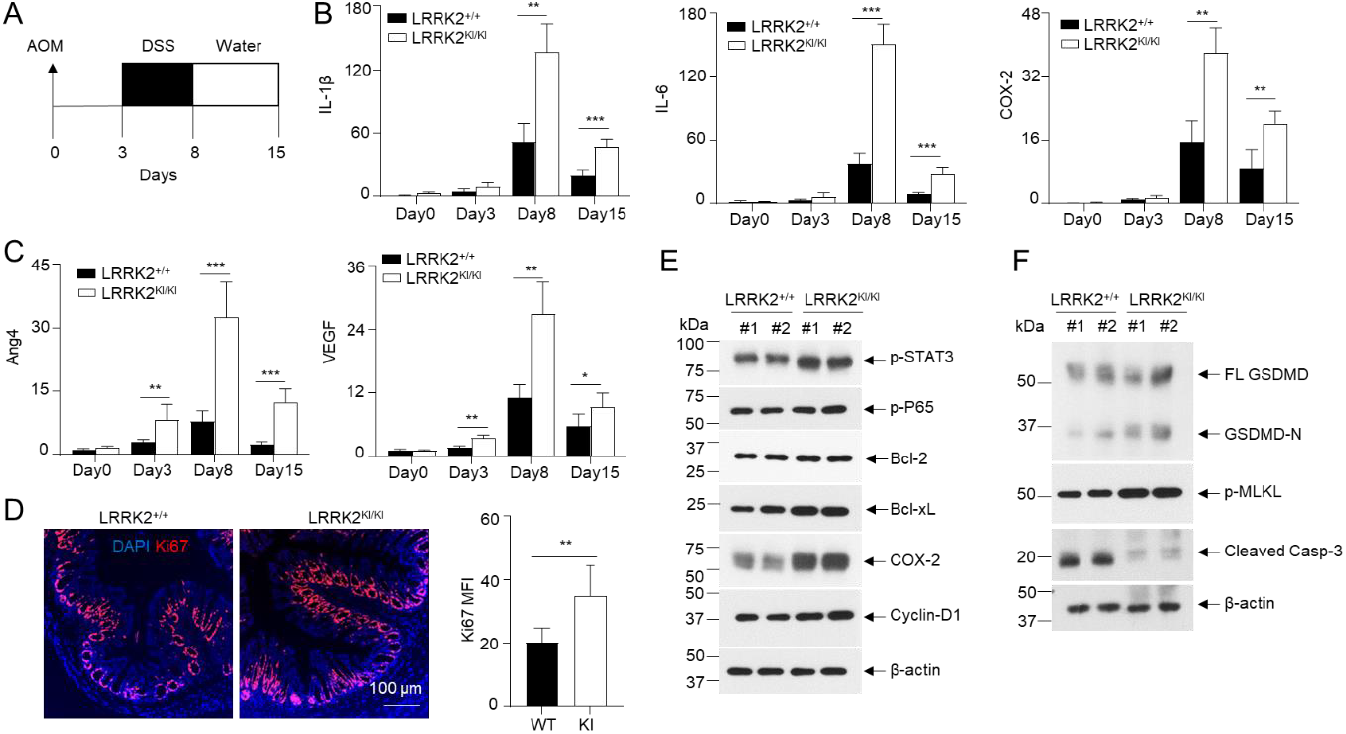
LRRK2 G2019S promotes inflammation and cell proliferation during early stage of tumorigenesis. (A) Schematic timeline for AOM and DSS treatment. (B) Real-time PCR of kinetic inflammatory gene expression in colon tissues from LRRK2^+/+^ and LRRK2^KI/KI^ mice as indicated. (C) Kinetic gene expression of Ang4 and VEGF in colon tissues from LRRK2^+/+^ and LRRK2^KI/KI^ mice were measured by Real-time PCR. (D) Ki-67 staining of colon tissues from LRRK2^+/+^ (WT) and LRRK2^KI/KI^ (KI) mice 15 days after treatment. Mean florescence intensity (MFI) was quantified by Image J. (E) Immunoblot analysis of key proteins involved in colon tumorigenesis as indicated. Lysates were prepared from whole colon tissues of LRRK2^+/+^ and LRRK2^KI/KI^ mice 15 days after treatment. Numbers represent individual mouse from each group. (F) Immunoblot analysis of markers of cell death from colon epithelial cells of LRRK2^+/+^ and LRRK2^KI/KI^ mice 15 days after treatment. n=3 mice/group, p values were determined by Student’s t-test. Data represent mean ± SD. *p<0.05, **p<0.01 and ***p<0.001. Data are representative of two independent experiments.

In the AOM/DSS model of CAC, AOM introduces mutation and genetic instability in the gut epithelium and DSS promotes inflammation in the tumor microenvironment, together they induce robust colon tumorigenesis (39,40). Consistent with this, LRRK2 G2019S mice did not development colon tumors when the mice were subjected to single AOM injection or repeated DSS treatment alone 65 days after treatment (data not shown). Consistent with the hyper expression of IL-1β and IL-6 in the colon tissues of LRRK2 KI mice (**Fig. 6B**), immunoblot analysis demonstrated increased p-STAT3 and p-P65 levels in the colon tissues of LRRK2 KI mice compared to those in WT controls (**Fig. 6E**). Correspondingly, we observed elevated levels of COX-2 and Cyclin-D1 in the colon tissues of LRRK2 KI mice (**Fig. 6E**). In the DSS-induced acute colitis model, we observed increased necrosis in the epithelium of LRRK2 KI mice compared with WT controls (**Fig. 5B** and **Fig. 5D**). We then asked if LRRK2 G2019S promotes necrosis in the gut epithelium at the early stage of colon tumorigenesis. Immunoblot analysis revealed higher levels of GSDMD-N and p-MLKL in the IECs from LRRK2 KI mice early after AOM/DSS treatment (**Fig. 6F**), suggesting increased necrosis. In contrast, we observed decreased cleaved caspase-3 in the IECs from LRRK2 KI mice, which may be explained by the results that hyperactivation of NF-κB and STAT3 in LRRK2 G2019S IECs upregulated the levels of Bcl-2 and Bcl-xL (**Fig. 6E**), which play critical roles in the anti-apoptosis response. Therefore, our data suggested that the overall impact of LRRK2 G2019S is to promote IEC proliferation at the early stage of colon tumorigenesis despite the fact that it can also promote necrosis. Necrosis promoted by LRRK2 G2019S may play a critical role in enhancing pro-tumorigenic inflammation.

## Discussion

While several studies have shown that PD patients who carry LRRK2 G2019S mutation have increased risks of developing cancers including colorectal cancer (21,22), the underlying mechanisms remain largely unknown. Here we demonstrate that LRRK2 G2019S promotes colon tumorigenesis in a mouse model of CAC. With the genetic tool of LRRK2 G2019S KI mice, we further uncovered that LRRK2 G2019S promotes intestinal inflammation in the DSS-induced colitis model. Considering the well-established positive correlation between colitis and colorectal cancer, our data suggest that LRRK2 G2019S-mediated colitis potentially promotes the pathogenesis of colorectal cancer in LRRK2 G2019S PD patients.

Though LRRK2 has been identified as the gene most associated with PD, meta-GWAS has also unraveled LRRK2 as a major susceptibility gene for Crohn’s disease (CD)(11-13). Of note, recent exome-sequencing analyses further revealed novel LRRK2 variants shared by both PD and CD including LRRK2 N2081D (36). LRRK2 G2019S is the most common genetic determinant of PD which is located in the kinase domain. Intriguingly, the N2018D variant is located in the same domain of G2019S. This promoted us to speculate that LRRK2 G2019S might promote intestinal inflammation and colitis. Using the DSS-induced colitis model, several groups have tested the role of LRRK2 in intestinal inflammation. Using LRRK2-deficient mice, Liu et al found that LRRK2 ablation promotes colitis in the DSS-induced acute colitis model (57). In contrast with this, a later study by another group suggested that high expression of LRRK2 by using LRRK2 transgenic mice also promotes intestinal inflammation using the same animal model and LRRK2 kinase inhibitor attenuates colitis severity in both LRRK2 transgenic mice and WT controls (41). Consistent with the latter study, Lin et al used LRRK2 G2019S transgenic mice and found that LRRK2 G2019S promotes intestinal inflammation in a DSS-induced chronic colitis model by upregulation of TLRs, NF-κB, and proinflammatory cytokines, especially TNF-α (58). One disadvantage of using transgenic mice is that the site of integration of the transgene into the genome can seriously affect tissue specificity and levels of transgene expression (59). Therefore, we used LRRK2 G2019S KI mice to overcome this drawback and revealed that LRRK2 G2019S KI mice exhibited significant weight loss, shortened colon length, increased spleen weight, and heightened gut damage compared to WT littermates, suggesting LRRK2 G2019S promotes colitis in the DSS-induced acute colitis model. In addition, inhibition of kinase activity of LRRK2 attenuated colitis severity in both KI mice and WT controls, suggesting kinase activity of LRRK2 plays a crucial role in intestinal inflammation. This result is consistent with the observation from a recent report where the research used DSS as a trigger to induce chronic intestinal inflammation in LRRK2 G2019S KI mice to promote α-synuclein-induced parkinsonism in this strain (60). They found that LRRK2 G2019S promotes DSS-induced chronic gut inflammation. Of note, the approaches used to induce colitis between this group and our group are different, but our results from G2019S KI mice together with the previous two studies using LRRK2 (41) or LRRK2 G2019S transgenic mice (58) support the notion that gain-of-kinase activities of LRRK2 promotes intestinal inflammation.

Though several studies as stated above have implicated LRRK2 kinase activity plays a critical role in DSS-induced acute and chronic colitis, the underlying mechanisms remain elusive. Inflammasomes play essential roles in gut homeostasis and intestinal disorders including IBD (52). Though research on the roles of inflammasome activation and its downstream effectors in experimental colitis models, such as the DSS-induced colitis model, have not obtained very consistent results or even opposite results sometimes, the clinical studies, however, suggest a positive correlation between inflammasome hyperactivation and colitis, especially IBD (52,61). Gene polymorphisms of NLRP3 and IL-18 has been suggested to confer IBD susceptibility (62-64). Mononuclear cells isolated from lamina propria of active colonic lesions in IBD patients produce higher levels of IL-1β and IL-18 (65-67), and colon IL-1β levels correlated with disease activity (68,69). Single-cell immune profiling revealed macrophages/monocytes from inflamed intestine tissues of IBD patients manifest IL-1β signatures (70). Furthermore, mutations in the IL-18R1-IL-18RAP locus are associated with susceptibility to IBD (11,71,72). However, the most direct evidence for the positive correlation between hyperactivation of inflammasome and IBD has come from the identification of autoinflammation with infantile enterocolitis (AIFEC). AIFEC is caused by inborn errors of NLRC4 which lead to hyperactivation of NLRC4 inflammasome and drive the pathogenesis of the disease (53,54). We previously discovered that LRRK2 is critical for the activation of NLRC4 inflammasome and its kinase activity is important for this function and host defense (8). In the current DSS-induced colitis model, we found that LRRK2 G2019S promotes inflammasome activation and subsequently leads to high production of IL-1β and IL-18 in the gut epithelium. Yet, we are not sure whether this is via NLRC4 inflammasome or others. This should be further addressed in future studies.

Upon inflammasome activation, another outcome is the cell pyroptosis in addition to producing mature IL-1β and IL-18. Pyroptosis is another form of programmed necrosis in addition to necroptosis (73). Consistent with the hyperactivation of the inflammasome in the gut epithelium of DSS-treated LRRK2 G2019S KI mice, we observed enhanced GSDMD cleavage, suggesting robust necrotic cell death in the intestinal epithelium. Furthermore, we also observed increased necroptosis. However, we did not see increased apoptosis, suggesting LRRK2 G2019S does not promote PANoptosis (74). Intriguingly, a recent study suggested that LRRK2 G2019S perturbs mitochondrial homeostasis and reprograms cell death pathways in macrophages by promoting cleaved GSDMD location on mitochondrial membrane(56). The outcome of this function is that G2019S variant promotes necroptosis instead of pyroptosis. We also found an increase location of GSDMD-N on the mitochondrial fraction of IECs from LRRK2 G2019S mice after DSS treatment, thus suggesting that LRRK2 G2019S not only promotes MLKL-mediated necroptosis in macrophages but also in epithelial cells. This result also suggests that the enhanced necroptosis in LRRK2 G2019S gut epithelium may be partially due to the translocation of GSDMD-N in the mitochondrial promoted by G2019S mutation. Consistent with the phenotype in LRRK2 G2019S macrophages, we also observed increased oxidative stress in the LRRK2 G2019S IECs as evidenced by the elevated production of ROS. As we know, ROS can promote inflammasome activation, thus potentially forming a vicious circle of inflammation in the intestine of LRRK2 G2019S KI mice. IEC death is one of the hallmarks of IBD. Elevated apoptosis and necroptosis of IECs are positively linked to the severity of inflammation in IBD (75-77). IEC death could lead to gut barrier dysfunction/erosion, microbial dysbiosis, and systemic spread of pathogens (78-80). In addition, necrotic cell death leads to the release of DAMPs, such as ATP, HMGB1, IL-1α, hyaluronan, and IL-33 et al (81), which would further promote intestinal inflammation. Collectively, our results suggest that LRRK2 G2019S promotes intestinal inflammation at multiple levels, and the relative contribution of these distinct pathways to colitis warrants further investigation.

It has been well established that IBD patients have an increased risk of developing colorectal cancer due to sustained intestinal inflammation (24,25,28,29). Inflammation promotes colon carcinogenesis at multiple levels and via various pathways (30). Inflammation-induced oxidative stress can initiate tumorigenesis via DNA damage and increased oxidative stress can inversely promotes inflammation(82). Inflammation can lead to impairment of the gut barrier and dysbiosis which will impact the colon tumorigenesis (83). Various inflammation-triggered signaling pathways are demonstrated to play crucial roles in colon tumorigenesis, especially NF-κB and STAT3 signaling (84-87). Our data suggested that LRRK2 G2019S promotes the production of IL-1β, IL-6 and IL-11 in colon tumor microenvironment, which activates the NF-κB and STAT3 signaling pathways. As such, we observed increased phosphorylation of STAT3 and p65 in LRRK2 G2019S colon tumors together with upregulation of the downstream molecules of Bcl-xL, Cyclin D1 and COX-2. However, how LRRK2 G2019S mediates these distinct signaling pathways and contributes to colon tumorigenesis remains to be explored in the future. In conclusion, our study provides important insights into the molecular mechanisms linking the LRRK2 G2019S mutation to colon carcinogenesis. We propose that LRRK2 G2019S promotes intestinal inflammation which creates an inflammatory microenvironment for the pathogenesis of colorectal cancer observed in LRRK2 G2019S PD patients. Targeting LRRK2 kinase activity may become a novel therapy for colon cancer patients, particularly in those with G2019S mutation or elevated LRRK2 expression.

## Authors’ Contributions

Y.W. designed and performed most of the experiments, interpreted the data, and wrote part of the manuscript. J.G. and T.S. aided in mouse genotyping, immunoblotting, and real time PCR. Y.X. instructed and performed pathology analysis. T.M., P.G. and S.S. contributed to data interpretation and manuscript revision. Z.K. was integral for experimental design, manuscript writing, data interpretation and project coordination.

## Acknowledgments

This work was supported by grants from national institution of health (NIH R21AG076895, R01 NS104164 to Z.K.).

