## Supplemental Table 1 and 2 for "LRRK2 G2019S promotes the development of colon cancer via modulating intestinal inflammation"

**Supplemental Table 1. Antibody list**

| Antibodies | Catalog number | Company |
| --- | --- | --- |
| p-STAT3 | 9145T | CST |
| P-P65 | 3033T | CST |
| P-Bcl-2 | 2827S | CST |
| Bcl-XL | 2764S | CST |
| COX-2 | 12282S | CST |
| Cyclin-D1 | 55506T cyclinD1 | CST |
| β-actin | 3700S | CST |
| IL-β | AF-401-NA | R&D |
| Caspase1 | AG-20B-0042-C100 | AdipoGen |
| GSDMD | ab209845 | Abcam |
| Caspase3 | 9661 | CST |
| Caspase8 | 8592S | CST |
| p-RIPK1 | 3493S | CST |
| p-MLKL | 91689S | CST |
| VDAC1 | 55259-1-AP | Proteintech |
| Ki-67 | 12202 | CST |

**Supplemental Table 2. Primer list**

| Primer name | Primer sequence |
| --- | --- |
| IL-1β F | ACTGTTTCTAATGCCTTCCC |
| IL-1β R | TGGTTTCTTGTGACCCTGA |
| IL-6 F | TAGTCCTTCCTACCCCAATTTCC |
| IL-6 R | TTGGTCCTTAGCCACTCCTTC |
| IL-11 F | CTGACGGAGATCACAGTCTGGA |
| IL-11 R | GGACATCAAGTCTACTCGAAGCC |
| IL-17 F | CCTCACACGAGGCACAAGTG |
| IL-17 R | CTCTCCCTGGACTCATGTTTGC |
| IL-22 F | GCTTGAGGTGTCCAACTTCCAG |
| IL-22 R | ACTCCTCGGAACAGTTTCTCCC |
| IL-23 F | CCTTCTCCGTTCCAAGATCCT |
| IL-23 R | ACTAAGGGCTCAGTCAGAGTTGCT |
| TNF F | GACCCCTTTACTCTGACCCC |
| TNF R | AGGCTCCAGTGAATTCGGAA |
| COX2 F | GCGACATACTCAAGCAGGAGCA |
| COX2 R | AGTGGTAACCGCTCAGGTGTTG |
| NOS-2 F | GAGACAGGGAAGTCTGAAGCAC |
| NOS-2 R | CCAGCAGTAGTTGCTCCTCTTC |
| CSF-1 F | GCCTCCTGTTCTACAAGTGGAAG |
| CSF-1 R | ACTGGCAGTTCCACCTGTCTGT |
| CCL7 F | CAGAAGGATCACCAGTAGTCGG |
| CCL7 R | ATAGCCTCCTCGACCCACTTCT |
| CXCL9 F | CAGAACCTCCCACGTAGCTTTC |
| CXCL9 R | GCTCTGAAGATGGGATCAAGTTAATA |
| GM-CSF F | GGCCTTGGAAGCATGTAGAGG |
| GM-CSF R | GGAGAACTCGTTAGAGACGACTT |
| Ang4 F | GGTTGTGATTCCTCCAACTCTG |
| Ang4 R | CTGAAGTTTTCTCCATAAGGGCT |
| VEGF F | CTGCTGTAACGATGAAGCCCTG |
| VEGF R | GCTGTAGGAAGCTCATCTCTCC |
| Wnt5a F | GGAACGAATCCACGCTAAGGGT |
| Wnt5a R | AGCACGTCTTGAGGCTACAGGA |
| MMP10 F | TGCTGCCTATGAGGCTCACAAC |
| MMP10 R | GGAGGAAAACCGAGAGTGTGGA |
